# Fifty Years After: The N1 Effect Travels Down to the Brainstem

**DOI:** 10.1101/2024.02.23.581747

**Authors:** Daniel J. Strauss, Farah I. Corona-Strauss, Adrian Mai, Steven A. Hillyard

## Abstract

Fifty years ago, it was reported that selective attention affects the N1 wave in auditory event–related potentials. We revisited the original study design but integrated the state of the art knowledge on short auditory stimuli and neural signal processing. In particular, one series of tone bursts has been replaced by chirp stimuli which are optimized to evoke consistent brainstem potentials at low and medium stimulation levels. Auditory selective attention affected the chirp– evoked response in subcortical structures, even at level of the inferior colliculi. A single–trial time–frequency analysis of the full–range (0–250ms) event–related potentials showed that selective attention increases the spectrotemporal consistency across trials in the corticofugal auditory pathway, at least from the N1 wave down to the auditory brainstem response.

---

When we focus on one sound source in a noisy environment, attention modulates the representation of that sound along the auditory pathway. The classic experiment to analyze this type of attention in humans is by event–related potentials, i.e., electrical potentials recorded from the scalp, while the participants have to listen selectively to pip sound sequences. Fifty years ago, it was reported that attended pip sounds enhanced the event–related potential 100ms after the stimulus, a cortical potential and an ‘early’ effect at that time. In the presented study, we updated this classic experiment by integrating chirp sounds. These sounds, which were already appreciated in the ancient Maya culture, evoke an enhanced neural response in very early structures of the auditory pathway. We report that attention enhances the event–related potential response in the brainstem as early as 5ms after the chirp presentation.

In a seminal paper in 1967, Näätänen argued convincingly that experiments aimed at studying physiological mechanisms of selective attention must be designed so as to rule out non–specific processes such as global arousal or alertness [1]. The key design feature he recommended was to present attended and ignored stimuli in unpredictable order to prevent different levels of arousal or other non-selective preparatory states from arising prior to the attended stimuli. With such randomization, a brain response that differed between attended and ignored stimuli could more confidently be interpreted as due to the selective processing (enhancement or suppression) and not to a non–selective arousal mechanism.

The first electrophysiological study of auditory attention in humans to incorporate this design feature was carried out by [2], who recorded event–related potentials (ERPs) to tone pips presented to the right and left ears in random order. The major finding was that the prominent N1 component of the auditory ERP with a latency of 60–150ms was enhanced in response to attended–ear tones. This “N1 effect” was interpreted as an early selection of attended channel inputs for further processing in the manner of a sensory gain control mechanism. An alternate hypothesis was put forward by [3], however, who proposed that the increased N1 amplitude with attention was not due to an increase in the evoked N1 component itself but rather to an endogenous “processing negativity” that was superimposed on the N1 and reflected the feature–specific analysis of the attended stimulus. Further studies (4; reviewed in 5) found that both mechanisms (N1 amplitude increase and added processing negativity) could contribute to the N1 effect, with the processing negativity onsetting later and lasting longer. It was also determined that the N1 component and its modulation with attention arose from neural generators in the auditory cortex [6, 7, 8]. An earlier cortical component in the mid-latency range (20–50ms) was also found to be modulated by attention prior the N1 effect [9, 8], lending support to the proposal that attention acts as a gain control over early evoked neural activity in auditory cortex.

What has remained unresolved in the 50 years since the N1 effect was reported is whether selective attention can modulate auditory input at subcortical levels. The wealth of descending pathways from cortex to different levels of the auditory brainstem pathways [10, 11, 12] could conceivably impose selective processing of attended stimuli in the brainstem that could be passed along to the auditory cortex. Numerous studies have investigated this possibility through scalp recordings of the auditory brainstem evoked response (ABR, waves I-VI) elicited within the first 10ms after sound (click or tone) presentation; the broad consensus of these studies has been that the ABR is invariant to manipulations of attention [13, 14, 15, 16, 17, 18, 19, 20, 21, 9]. More recent studies of the auditory frequency following response (FFR) to speech sounds, however, have revitalized this controversial question of whether attention can affect auditory transmission in the brainstem.

The FFR is a near–sinusoidal oscillatory potential in the range of 100–300Hz that originates from the auditory midbrain pathways and is phase–locked to the voicing frequency of the speech input. Considerable evidence has accumulated that the scalp– recorded FFR to speech sounds shows increased amplitude and/or phase–locking when those sounds are attended [22, 23, 24, 25].

It remains unclear, however, whether this top–down modulation of evoked neural activity in the brainstem is specific to the oscillatory FFR to speech stimuli or whether it affects a broader range of stimuli, see also [26]. It is also unclear how soon after stimulus onset the selective FFR modulation begins.

The present study reopens the question of whether the ABR can reveal attentional modulation of short latency evoked activity in the brainstem pathways by presenting tonal “chirp” stimuli in a dichotic listening design that paralleled that of the original [2] study. The chirp is a brief tone burst that is frequency modulated from low to high in such a way it activates the entire cochlea in synchrony and elicits ABRs at medium to low intensities [27, 28]. Participants were presented with randomized sequences of chirps to one ear and 800 Hz tone bursts to the other ear with instructions to attend to one ear at a time and report occasional targets of lower intensity in the attended ear. ERPs were recorded to all stimuli from a dense (128–channel) electrode array with the aim of examining attentional modulation of auditory input at brainstem, mid–latency, and long–latency cortical levels. Complementary to this multichannel configuration, we used an additional set of single electrodes to have same vertex configuration as in seminal study in [2].

## Results

### Spectrotemporal Attention Signatures

In Fig. 1, we have shown the full–range ERPs evoked by the chirps and tones for the attended and the ignored condition using the original EEG electrode configuration in [2]. For both types of stimuli, we see a clear N1 effect as described in [2], i.e., the negative deflection around 100ms is larger for the attended condition. As expected, the responses from earlier anatomical structures along the auditory pathway are much clearer represented for the chirp stimuli than for the tones. For instance, a P20–P50 attention effect [29] can be observed for the chirp. In particular, the prominent wave V in the ABR (around 5ms) which is anatomically associated with the inferior colliculi (ICs), see [30, 31], is clearly noticeable for the chirp stimuli. In our following discussions, we present only the results for the chirp stimuli whereas complementary analyses for the tones are given in the supporting information (SI). An additional source localization analysis using the multichannel data to the chirp stimulation confirmed the locus of wave V, see SI.

**Figure 1:**
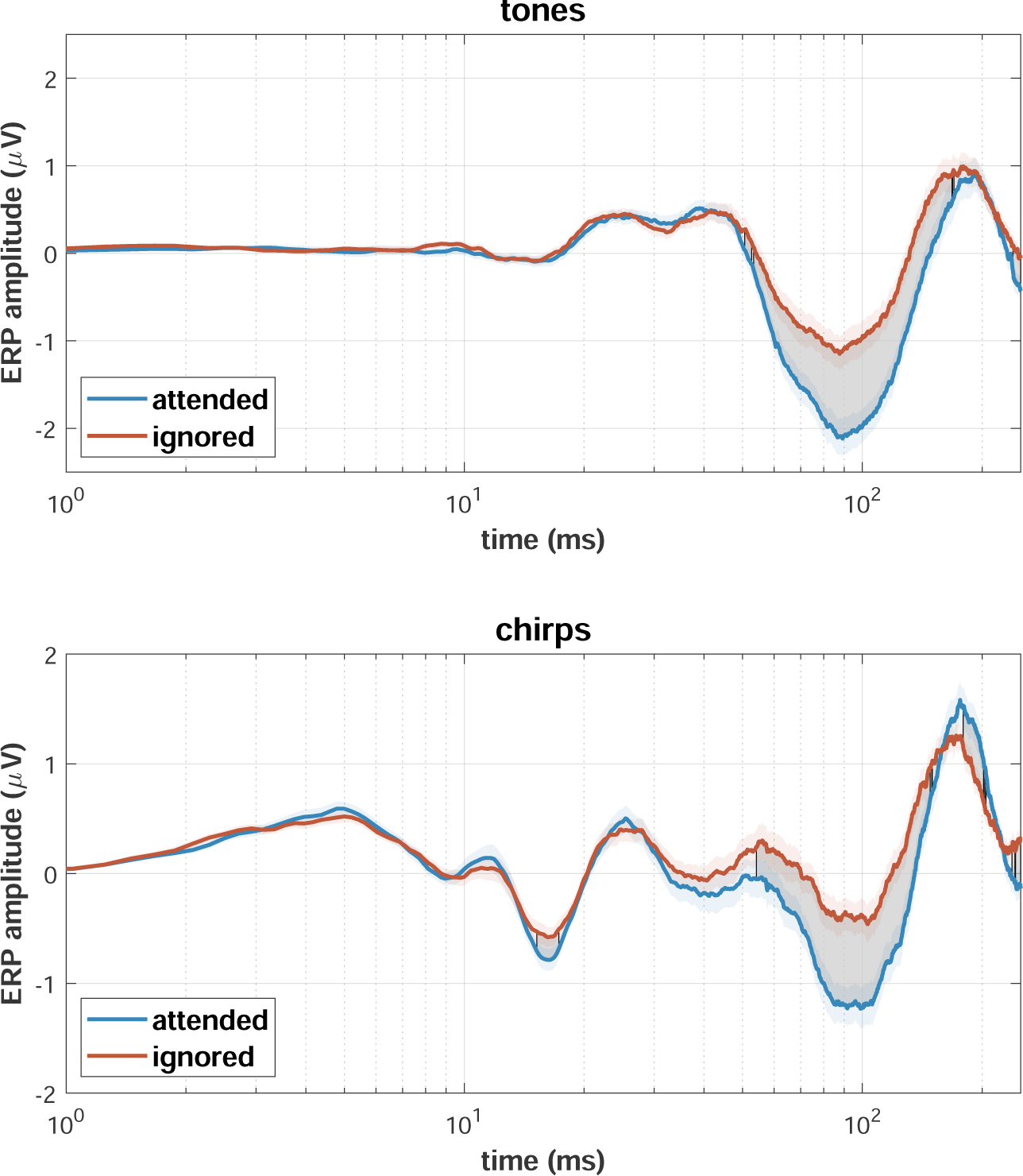
Grand average vertex configuration broadband ERPs to tones (top) and chirps (bottom) on a logarithmic time–scale. The colored shaded areas show the standard error across participants and gray areas highlight time periods which indicated a significant difference between conditions (*p* < 0.05; two–tailed within–participants t–test with false discovery rate (FDR) correction). While responses to both stimuli present a dominant “N1 effect” at around 100ms, the chirp–evoked ERPs additionally exhibit noticeable attentional modulations at multiple sites along the auditory pathway, even reaching significance for an early negative deflection at around 16.25ms.

As full–range ERPs cover several spectrotemporal scales [32, 33], we employed a time– frequency analysis to map frequency specific signatures of top–down attention in the corresponding single–trial sequences making up the averaged ERPs to the chirp stimuli, see Fig. 2. Even though similar difference patterns have been observed for the inter– trial phase coherence (ITPC) for the N1 wave, see [34, 35], there are several significance blobs in the time–frequency distribution of the ITPC well before 100ms.

**Figure 2:**
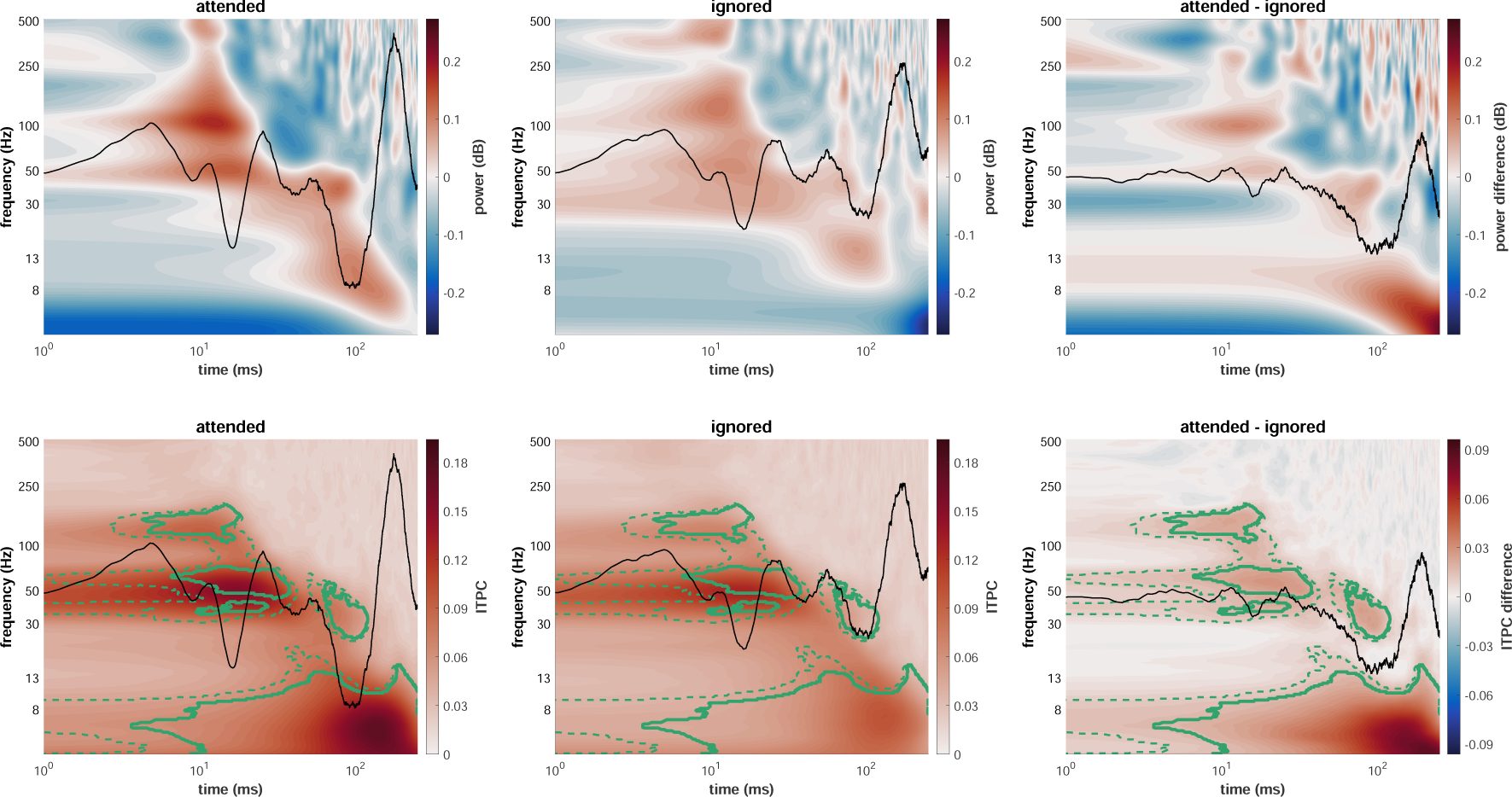
Wavelet power (top) and ITPC (bottom) analysis of vertex broadband single sweeps to attended (left) and ignored (middle) stimuli as well as the resulting difference map (right; attended - ignored). Black waveforms represent identically scaled condition–specific grand average ERPs (left and middle) and their difference wave (right). Time– frequency areas which indicated significant differences between conditions are represented by green lines (*p* < 0.05 for dashed lines and *p* < 0.01 for solid lines; two–tailed within–participants t–test with FDR correction). Note that while the ITPC significance mask shows clear structures matching the individual sub–components of the auditory full–range ERP, the power is absent of any significant patterns.

### Signs of Attention in the Human Brainstem Response

In particular, the interval centered around wave V in chirp–evoked ABRs is different for the attended and the ignored condition and thus shows a modulation by selective attention. The corresponding high–frequency band (105.3–185.0Hz) is in line with fibre tract models of ABRs [36] and the logarithmic up–chirp behavior of the ERPs, see SI. Derived from the conventional vertex configuration, we have shown the averaged time– domain ERP filtered in this frequency band along with its version using a conventional chirp–evoked ABR filter [37] in Fig. 3 (top). In Fig. 3 (middle), the first principle component from the conventionally filtered [37] array electrode configuration is shown. It is noticeable that this data, even though derived from different electrodes, represents the same ABR effect. To analyze this effect even further in the time–domain using a nonlinear method, we have shown the grand averages of the 4th mode of a 7–mode variational mode decomposition (VMD) of the early (*−*10–50ms) mean ERPs across participants in Fig. 3. The averaged principal modes of both conditions have nearly identical center frequencies (126.5Hz for attended, 126.1Hz for ignored) well within the ITPC significance blob of the wave V interval in Fig. 2 (bottom). Complemented by a time–frequency super–resolution analysis in the SI, Fig. 3 shows that the high– frequency oscillation in the wave V interval is an intrinsic, principal mode of the ERP and not a byproduct of later components due the Heisenberg’s uncertainty principle when using predefined time–frequency atoms in linear transforms. Even though the (time–domain) amplitude of ABRs is fragile as compared to the ITPC, see [28], the shown bandlimited intrinsic mode exhibits amplitude significant differences. A more detailed statistical analysis can be found in the SI.

**Figure 3:**
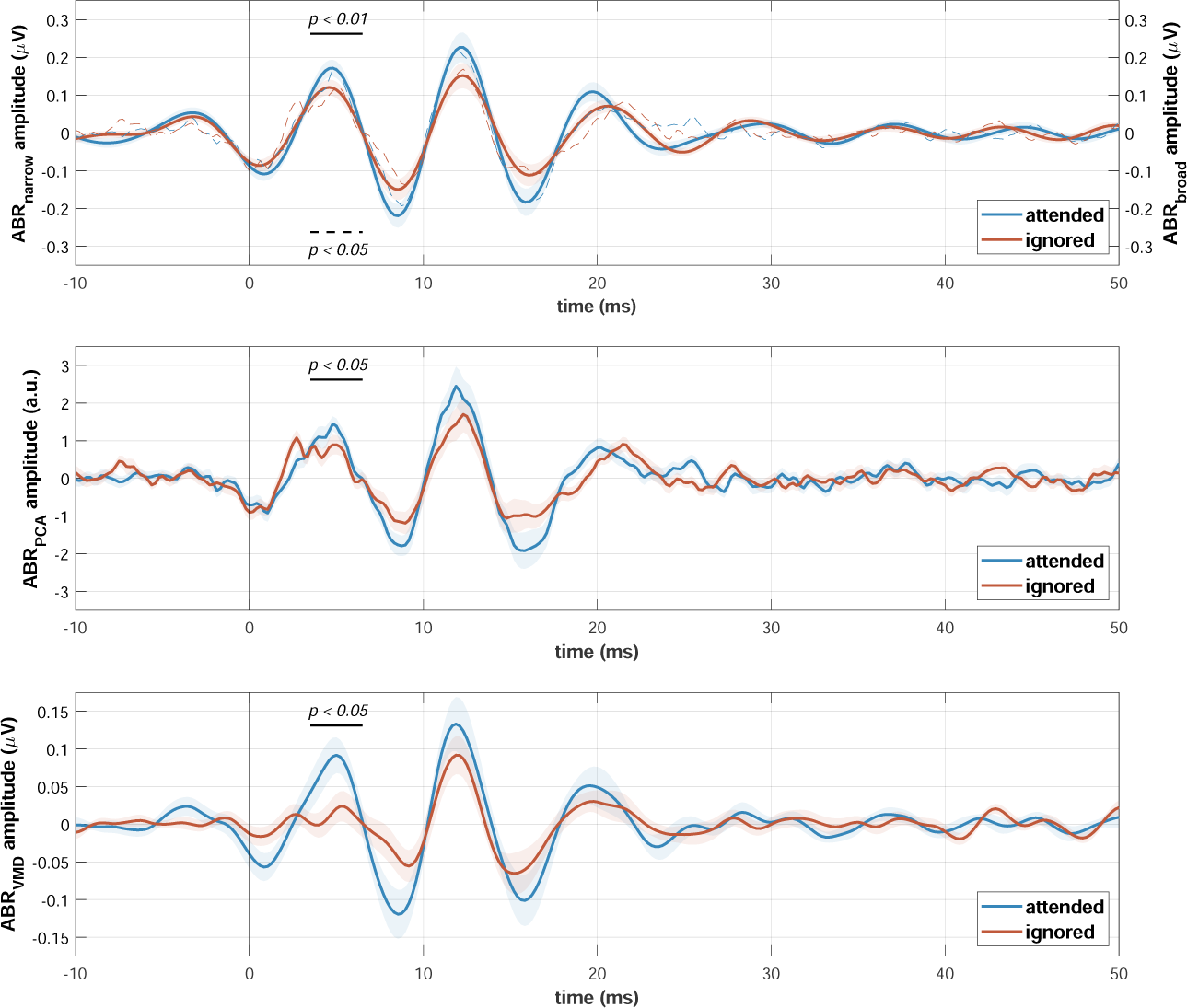
Grand average waveforms of enhanced ABR representations. ABRs from mean vertex ERPs were extracted via bandpass filtering (top) within a narrow (105.3–185.0Hz; left y–axis) and a broad (100–1500Hz; right y–axis) passband as well as a VMD analysis (bottom). ABRs in multi–channel recordings were enhanced by applying the identical broad bandpass filter followed by a PCA analysis (middle). The colored shaded areas show the standard error across participants. In addition to the highly similar high–frequency behavior within the first 20ms of the ABR, all four representations exhibit amplitudes in the wave V interval (highlighted by the horizontal black lines) that are significantly increased with attention (all *p* < 0.05; two–tailed within–participants t–test).

### Selective Attention induces Spectrotemporal Consistency

The attended chirps caused an increase in the ITPC for several time–frequency patches, see Fig. 2 (bottom), as well as an increase of the amplitude of several peaks and troughs in the averaged full–range ERP, see Fig. 1 and 3 but not in the energy of the individual trials, see Fig. 2 (top). Consistent with the direct modulation and increase in amplitude and energy related to the “N1 effect” in the averaged ERP, temporal precision in auditory processing has been linked to the attentional gain theory and an inter–trial correlation in the time–domain, see [38] who used the term temporal precision to map the temporal consistency across trials. In the following we use the term spectrotemporal consistency across trials (SCAT) to a generalize the term ‘temporal precision’ and refer to neural synchronization stability across trials in time and/or frequency, similarly to [39, 40].

Complementary to the ITPC analysis in Fig. 2 (bottom), Fig. 4 shows the mean correlation of the averaged ABR, AMLR, and ALR to the individual trials in the time-domain for the attended and the ignored condition (top: left plots) as well as the mean energy of the individual trials (top: right plots). It is noticeable that, in contrast to the energy, the correlation is significantly different between the attended and the ignored condition. Along with the ITPC analysis in Fig. 2 (bottom), this demonstrates that selective attention induces SCAT, i.e., a time–locked morphological stability of the individual responses in ABRs, AMLRs, and ALRs.

**Figure 4:**
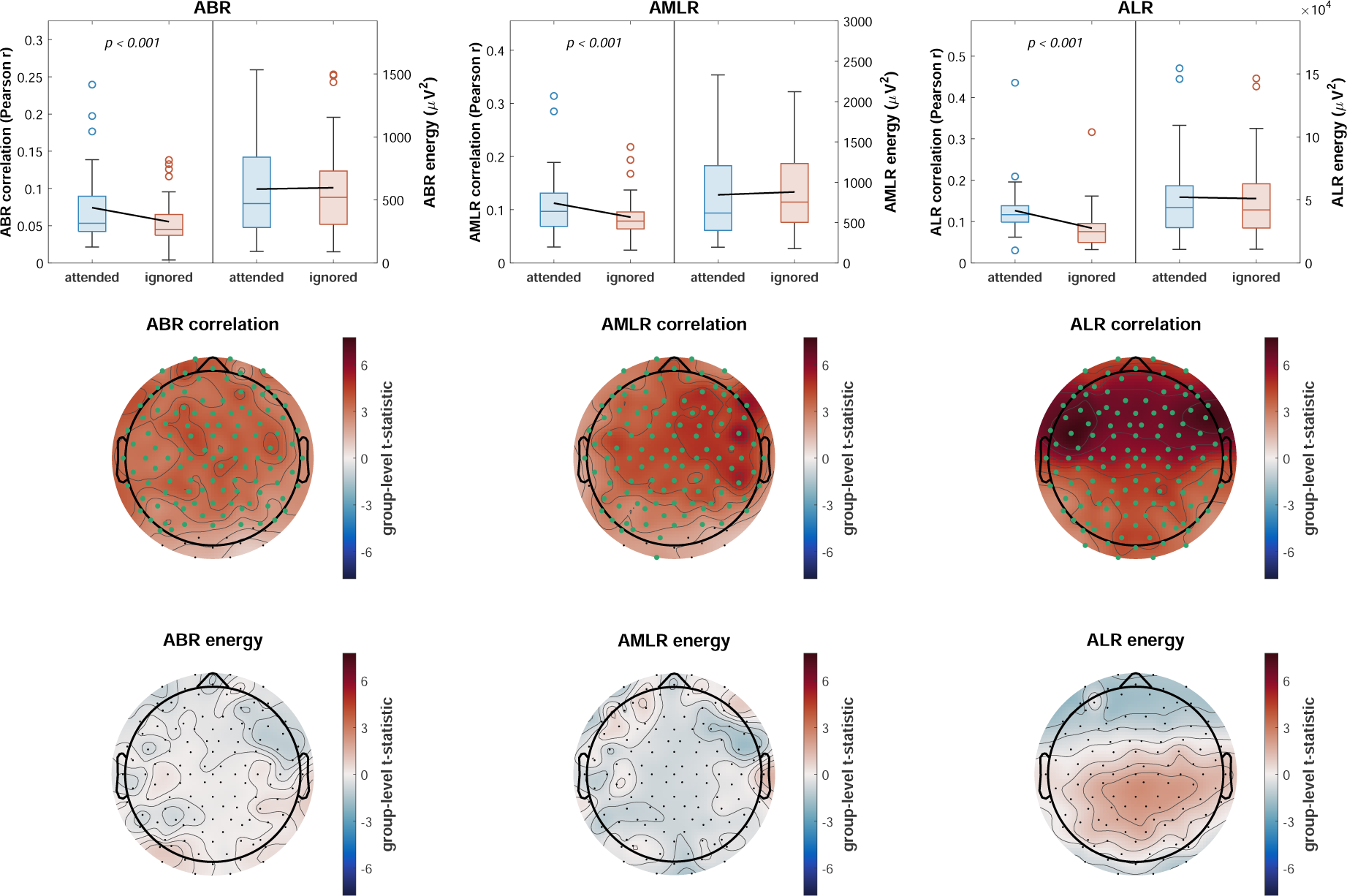
Time–domain correlation and energy analysis for the individual full–range sub–components (ABR, AMLR, ALR; left to right). Correlations were computed between the mean waveforms and each of its single responses, energy within each single response, and the results were averaged across sweeps. The figure presents the raw values for the vertex recordings (top) for correlation (left y–axis) and energy (right y–axis) and the group–level t–statistic topographies for multi–channel recordings for correlation (middle) and energy (bottom). Channels which indicated significant differences between conditions are marked by green dots. While the energy did not present any significant effects, the correlation was significant for all contrasts of vertex recordings (all *p* < 0.001; two–tailed within–participants t–test) and the vast majority of channels in all contrasts of multi–channel recordings (*p* < 0.05; two–tailed within–participants t–test with FDR correction).

To analyze the effect of SCAT in single–trials topologically, we depict the group-level t-statistic topographies for the high–res EEG recording (attended vs. ignored) for the mean time–domain correlation of the averaged ABRs, AMLRs, and ALRs to their single sweeps Fig. 4 (middle) and the mean energy of the responses across single–trials in Fig. 4 (bottom). It is noticeable that SCAT is discriminating between the attended and the ignored stimuli for all ERP types at almost all channels. The differences for the ABR are larger in central areas, lateralized towards the right hemisphere for AMLRs, and concentrated on frontotemporal regions for the ALRs. Regarding the processing locus, this wide–spread neural synchrony suggests a far–field influence in the case of ABRs, and more coordinated activity in a wide–spread circuitry for AMLRs, and in particular for ALRs. The mean energy of the individual single–trials does not exhibit differences and does also not follow the topological pattern of the SCAT measure. However, it is important to note that the result in Fig. 4 is specific for the respective potential (i.e., ABR, AMLR, and ALR) but not its individual temporal features (or waves), i.e., if several peaks and toughs of one potential change their energy in an opposing fashion, the net energy might remain the same.

## Discussion

Similarly to the “N1 effect” described in [2], we have shown that selective attention modulates auditory processing at much earlier stages of the auditory pathway for non-speech stimuli. As in the classic selective attention study in [2], we followed [1] and presented the attended and ignored stimuli in unpredictable order to prevent different levels of arousal or other non–selective preparatory states from arising prior to the attended stimuli, discriminating this approach from attempts using the FFR to speech stimuli with their inherent structure, see [26]. For this, we replaced one tone stimulus sequence in [2] by chirps which provide a more synchronous discharge of the entire cochlea and thus results in more structure in the full–range ERP.

The ABR interval of this ERP showed an increased amplitude in the averaged potential across trials in the interval of the wave V which is anatomically linked to the ICs [41, 42]. The ICs and subcollicular are targeted by massive efferent corticollicular projections [11, 12] and have been linked to attentional modulations in humans [43, 44]. Literature on later components in ABRs is less conclusive but linked these components to the activity of medial geniculate body (MGB) of the thalamus and auditory radiations [30, 42]. Using intracerebral recordings in human, [42] reported that the earliest activity in the auditory cortex was found at 17ms for tone burst stimuli. Earlier studies linked SCAT of the N1 wave assessed in time [38] or by the instantaneous phase relations spectrotemporally [34, 35] to the “N1 effect” of attention. Our study showed that this (macroscropic) SCAT is a general phenomenon of auditory selective attention, from the auditory cortex down to the brainstem. It does not just result in a high spectrotemporal consistency and precision of the full–range ERP peaks and troughs, but also in time– locked and phase–locked relations between them. Even though the link between the power of the averaged ERP, the power of individual sweeps and the ITPC as well as the link between amplitude variations and the instantaneous phase in the EPRs analysis are well understood meanwhile, see [45] and [46], respectively, it is difficult to derive any conclusion about the generative mechanism of ERPs from it due the gap between spatiotemporal scales linking macroscopic scales and microscopic generative phenomena [34, 47]. Moreover, the statistical analyses in Fig. 2 originated from two different data types, linearly scaled amplitude data and circular phase data. It is known that these data types have a different noise sensitivity in ERP analysis [48, 49]. Thus, due to the different scaling nature of the underlying data, these results merely reflect the sensitivity of detecting the effect using this particular data type and not the nature of the effect itself.

Studies in vision and mostly for corticocortical pathways suggest that neural synchrony might reflect the basic computational principle of top–down attention in a biased competition framework [50], see [51, 52]. In general terms and on the microscopic scale, this refers to the synchronization of the rhythmic response of neurons that are tuned to the spatial and featural attributes of the sensory input [51] or causal phase relations across efferent (downstream) brain structures in local neural circuits for top–down attention [52]. In fact, gain control by attention as fundamental principle [53] (Chapter 11) resulting in better neural synchrony or SCAT could stem from various microscopic phenomena, see [53, 54]. For instance, the basic circuitry to implement the normalization and inhibition mechanisms reviewed in [54], seems also to be present in corticollicular top–down pathways, see [55]. Due to the gap between spatiotemporal scales linking spikes and local field potentials to macroscopic ERPs [34, 47], it remains elusive to relate these macroscopic and microscopic synchronization phenomena in corticofugal modulation directly.

Generative models of ERPs, in particular, the evoked vs. the phase-resetting model, are discussed in literature since decades, e.g., [56, 57, 58, 59]. However, the inference of the underlying generative mechanisms from scalp ERP data represents mathematically an ill-posed inverse problem, see [60] for a detailed discussion. One conclusion from this is that, as scalp ERP data just gives access to the macroscopic effect such as SCAT and not the meso- and microscopic neuronal cause of the effect, an infinite number of models can be found using the one or the other generative model. Without constraining the space of possible solutions to valid generative models by specific onsite (invasive) measurements and microscopic data, a reasonable approximate solution to this inverse problem is elusive [60]. Therefore, it is beyond the scope of this paper to suggest generative models, let alone suggesting conjectures for generative models of the observed attention effects in the full–range ERPs. However, even though ABRs are known to increase the ITPC for higher frequencies (*>* 100Hz), see [28], what our study adds here is a very early (a few milliseconds) brainstem effect of attention regarding SCAT (as assessed by the averaged ERP amplitude, its ITPC, as well as its correlative precision in time). The corresponding high–frequency neural correlate is in line with fibre tract models of ABRs [36] and the logarithmic up–chirp behavior of the ERPs, see SI. Generative models which cover multiple frequency intervals, e.g., see [58], might integrate these results in future modeling efforts.

Even though the precise generative mechanisms of this ABR effect are still unknown, its manifestation of an enhanced wave V of the averaged ERP resamples the N1 effect reported as early sign of attention 50 years ago using a similar but carefully updated experimental paradigm.

## Methods

### Stimuli and Tasks

The auditory paradigm was constructed as closed as possible as in [2], but with some modifications explained in the following. The one schedule configuration remained having different sequences of stimuli in each ear, but we used broadband chirps in one ear and 800 Hz tone bursts in the other ear, see Fig. S1 in the SI. Each sequence had its own target stimulation, which was the same stimuli but 15dB softer in intensity. The stimuli were created using a sampling rate of 44.1kHz.

The tone burst lasted 50ms including rising and falling time of 12ms each, both taken from the 2 halves of a Hanning window of 24ms of duration. The chirp (see [27] and [28]) is the A-Chirp for 40dB SPL, with a frequency band of 100-9800Hz, resulting in a final duration of 14.9ms. Other chirps tested during previous piloting phase, particularly shorter in duration, did not evoke streaming perception in selective listening with the competing tones, even at faster stimulation rates. Chirps of shorter duration were more heard like a click sound and participants did not perceive a tonal structure, loosing their tonal capabilities for the detection task and particularly for maintaining auditory streaming. Chirps were chosen because of their broadband frequency nature, stimulating a larger portion of the cochlea even at medium and softer intensity levels, resulting in larger potentials [27]. In [61], results using modified versions of the the A-Chirp are presented, here the parameters of the chirps were modified so their final duration deviated approximately 15% from the original A-Chirp. In [62] found that changing parameters of the chirps, like stiffness constant of the basilar membrane, had a small effect on the resulting evoked response for the same subject. Additionally, [27] used a chirp based on otoacoustic emissions data, which had a duration of around 14ms. Authors concluded that although both chirps differed considerably in length, the resulting amplitude of the evoked potential hardly differ from one another for each subject tested.

The auditory paradigm was then generated using an inter stimulus interval (ISI) ranging from 0.25-0.4sec. The targets had a probability of appearance of 9.5% appearing randomly interposed every 3 to 20 stimuli. A duration of around 3min was calculated, i.e., around 300 stimulation per ear. Thus, in order to collect enough individual trials, epochs or sweeps, at least 20 repetitions of the auditory paradigm were necessary. In order to avoid participants to get familiar with only one stimulation file, ten conditions using the very same parameters, like the one described above were generated, resulting in different target positions but similar degree of difficulty regarding ISI, probability of appearance, etc.

### Participants and Electrophysiological Recording

Thirty one normal hearing young volunteers from the university environment participated in the experiments (26.8 *±* 3.7y, 23m/8f). After explanation of the experiment workflow, hearing sensitivity was measured (mean Pure Tone Audiograms (PTA) *<* 15dB SPL). After collecting the hearing thresholds for the chirp and tone burst (0dB Sensation Level (SL)), the stimulation was set to 40dB SL. Stimulation was presented via circumaural headphones (HDA300, Sennheiser, Germany).

A loudness fine tuning process followed by asking the participants to grade the loudness perception of both stimuli in both ears (chirps on right ear/ tone bursts on left ear and vice versa), assuring that in both cases, their loudness perception of both stimuli was the same. A maximal adjustment of *±* 3 dB was possible during this process. The resulting normal Hearing Level (nHL), i.e., the average 40dB SL of the stimuli were 71dB peSPL (59.5dB SPL) for the chirps and 48.6dB peSPL (52.1dB SPL) for the tone bursts. The average pe SPL for the mean 0dB SL for the chirp was 31.94dB (32.30dB SPL and 31.58dB SPL for the right and left ear, respectively, which are similar to the values reported in [27]), and 7.42dB (7.51dB SPL and 7.33dB SPL for the right and left ear, respectively) for the tone burst.

After a short training (2-3 min solving the task for chirps and ton bursts), participants learned to identify the target stimulation by pressing a button. Next, participants were sat in a comfortable armchair, instructed to move as less as possible during measurement times. A fixation ball was placed and was adjusted in front of the participants at sight level and 2m distance.

Skin preparation and electrode placement took place next, by attaching 9 Silver/Silver-Chloride (Ag/AgCl) passive electrodes on the upper and lower mastoids, vertex and one pair above each post auricular muscle (PAM). The PAM activity was monitored in order to rule out its recently described dependence on the direction of attention, see [63]. Subsequently, an EEG cap with 128 active sintered Ag/AgCl electrodes was placed (g.Scarabeo, gtec, Austria), adjusted and fulfilled with gel. Impedance were controlled to be lower than 10kΩ for the passive and 50kΩ for the active electrodes.

The electroencephalography (EEG) activity from the 133 electrodes, electromyography (EMG) activity of 4 electrodes, the corresponding stimulation trigger signals, and the signal of the button were acquired with a biosignal amplifier (gHIAMP, gtec, Austria) using a sampling rate of 9600Hz. The auditory stimulation and trigger signals were presented using a cross platform DAW (StudioOne, PreSonus, USA) coupled to a sound card (Scarlett 18i20, Focusrite,United Kingdom). The button and trigger signals, were conditioned before acquisition using a conditioner box (gTRIGBOX, gtec, Austria).

A set of 20 trials was acquired attending 10 times to the chirps and 10 times to the tone bursts. During the first 10 conditions, participants had either chirps on the left and tone bursts on the right or vice versa and in a pseudo-randomized order they solved 5 times the paradigm detecting the targets of the chirps and likewise 5 times for the tone. For the next 10 trials, the stimulation switched ears and the procedure was repeated. The ear at which the participants had the chirps the first ten measurements was also pseudo-randomized, in order to have around 50 percent of the participants starting with chirps on the right and the other 50 percent starting with chirps on the left. The latest was done with the idea for compensating for better ear or ear preference. After each trial short breaks were allowed, and participants were able to rest, eat small snacks and drink some water. Electrodes impedance were also controlled during these breaks. The stimulation files and further processing was done using software for scientific computing (Matlab and Simulink, The Mathworks, USA).

### Behavioral Data

After participants’ debriefing, it was found that 35.48% of the participants preferred or found it easier to solve the behavioral task for the chirps, 32.26% for the tones, 19.35% had no preferences, and 12.90% had a better–ear preference. In order to objectively evaluate their discrimination performance between standard and target stimuli, Matthews correlation coefficient (MCC) [64] was computed for each trial. Herefore, button presses were only classified as hits if the reaction occurred between 10–1500ms after target onset, and misses otherwise. While the mean MCC across participants and trials was slightly higher for chirps (0.82 *±* 0.10) compared to tones (0.80 *±* 0.10), a within– participants t–test (two–tailed) attested that the stimulus type had no significant effect on task performance (*t*(30) = 1.36, *p* = 0.19).

### EEG Preprocessing and Segmentation

EEG preprocessing was conducted in two stages with an initial stage for common processing of vertex and multi–channel recordings, and a final stage for artifact correction in the latter. As the earlobe electrodes were excluded from analysis and the positioning of two electrodes varied across participants, all following multi–channel procedures included 124 channels.

The study paradigm was designed to reliably evoke auditory full–range ERPs including the ABR, the AMLR, and the ALR. Since ABRs are preferably investigated using a mastoid reference ipsilateral to stimulation [32], all recordings were referenced to the mastoid ipsilateral to attended as well as ignored stimuli which resulted in two data sets per trial. Data were then decimated to 4800Hz, zero–phase comb–filtered at 50Hz and its harmonics up to 2400Hz, zero–phase bandpass–filtered in [1, 1500]Hz, and corrected for their DC–offsets. In the second stage, multi–channel recordings from all trials were concatenated (including both ipsilaterally referenced data sets) and the mean Pearson correlation between each electrode and its five nearest neighbors was computed. Channels presenting correlations below two standard deviations from the mean across all channels were interpolated using EEGLAB’s [65] spherical interpolation algorithm (6.52 *±* 1.90 per participant). In general, artifact correction was kept to a minimum to avoid removal of potentially subtle effects, as could be the case for common multi–channel techniques such as cleaning via independent component analysis. Data were then split into the individual data sets and centered around 0*µ*V.

ERPs for vertex recordings were extracted by segmenting the ongoing EEG from *−*1000 to 1000ms relative to onsets of standard stimuli exclusively to exclude target detection ERP correlates such as the P300. Since the stimuli differed considerably in their physical properties, true ERP onsets within the extracted segments were afterwards adjusted to stimulus–specific reference time points. These were specified as the rise time offset at 12ms for tones and stimulus offset at 14.9ms for chirps and served as their respective *t*_0_. To remove signal offsets while considering the latencies of the earliest ERP components, single sweeps were baseline–corrected by subtracting the mean potential between *−*2 and 2ms relative to adjusted ERP onsets. Finally, for each combination of stimulus (tones and chirps), condition (attended and ignored), and location (left and right), the first 1000 sweeps which did not exceed an absolute amplitude threshold of 100*µ*V were selected and pooled across locations for further analysis. The corresponding responses were extracted for multi–channel recordings and baseline–corrected as reported above. Since the grand average waveforms of vertex recordings only revealed a clear full–range structure for ERPs to chirps but not for tones (see Fig. 1), all following analyses were based on chirp–evoked responses if not stated otherwise.

### ERP Wavelet Analysis

Single sweeps from vertex recordings were mapped to time–frequency representations by means of continuous wavelet transforms with analytic Morse wavelets [66]. Specifically, data were analyzed at logarithmically spaced wavelet peak frequencies between 4 and 512Hz with 32 scales per octave. Because of this broad spectral range, the number of wavelet cycles within the central time–domain power window was linearly increased in [0.75, 2 (0.18 cycles increase per octave) to simultaneously provide satisfactory temporal resolution at low frequencies and spectral resolution at high frequencies, e.g., see [67]. This was achieved by fixing the wavelet family parameter *γ* at 3 and varying the order *β* between 1.85 and 13.16. The complex wavelet coefficients were then used to extract power– and phase–based time–frequency measures. Total power was computed by squaring the absolute values of the coefficients and averaging the resulting power matrices across sweeps. The final power matrices were then baseline– normalized independently for each wavelet scale by dividing the power at each time point by the mean power between *−*500 to *−*125ms and log–transformed (10*log*_10_) to decibels. Furthermore, ITPC was computed by extracting the instantaneous phase angle at each time–frequency point and calculating the mean resultant vector length [68] across sweeps in a point–wise manner. The corresponding analysis for the tone–evoked responses is provided in the SI.

### ERP Time–Domain Correlation and Energy Analysis

Contrasting ITPC between conditions revealed significant (*p* < 0.01) time–frequency clusters which corresponded to the individual full–range ERP sub–components (see Fig. 2). Projecting the significance masks onto the frequency axis, this resulted in frequency ranges of [105.3, 185.0]Hz (ABR), [44.3, 71.3]Hz (AMLR), and [4.0, 15.7]Hz (ALR). Following up on this result, vertex and multi–channel mean ERPs and their individual single sweeps were decomposed into the narrowband components ABR_narrow_, AMLR_narrow_, and ALR_narrow_ by applying zero–phase bandpass filters with the aforementioned passbands and extracting the signals within [0, 20]ms, [10, 50]ms and [40, 250]ms, respectively. For each sub–component, the correlation between the mean waveform and each of its individual sweeps as well as the energy (squared *ℓ*^2^-norm) of each single sweep were computed and averaged across sweeps.

### ABR Enhancement and Wave V Analysis

To enhance the ABR in a similar vein to the narrowband filtering procedure described above, ERPs were subjected to three additional types of time–domain analyses. The first approach employed a zero-phase bandpass filter with a conventional broad chirp– evoked ABR passband of [100, 1500]Hz [28] which was applied to the mean ERPs of vertex recordings to obtain broadband ABRs (ABR_broad_).

For the second analysis, mean ERPs from multi–channel data were filtered analogous to ABR_broad_ and submitted to a principal component analysis (PCA). This allowed to exploit the rich information provided by the high–density recordings for separating the ERPs via a spatial filter into components with varying degrees of contribution to the overall neural activity across the scalp and provided a denoising effect for the previously minimally cleaned data. PCA was carried out over the latency range [*−*10, 50]ms to maximize sensitivity towards the ABR, and the ABR component (ABR_PCA_) was identified as the component which accounted for most of the variance in the sensor recordings.

To analyze the high–frequency behavior of the ABR without any fixed a priori specifications about the spectral content of the intrinsic activity, mean ERPs from vertex recordings were decomposed into seven intrinsic modes by applying a variational mode decomposition (VMD) using the alternate direction method of multipliers [69]. As the VMD employs an optimization of the Hardy space representation of the modes in terms of their *H*^1^–Sobolev regularity for spectral compaction in the Fourier domain and the auditory full–range ERP follows the spectrotemporal trend of an up–chirp behavior (see SI), the analysis was again restricted to the time interval of [*−*10, 50]ms to keep the degree of non–stationarity low. The center frequencies of the modes were initialized as the peaks of the ERP’s Fourier spectrum and the minimization of the augmented Lagrangian functional was performed using a regularization constant of *α* = 1000 and a maximum number of 10000 iterations. The intrinsic ABR activity was detected by averaging the resulting modes across participants and analyzing them in the frequency– domain. Since only the fourth mode was concentrated well within the ABR_narrow_ passband with nearly identical peak frequencies of 126.5Hz for the attended and 126.1Hz for the ignored condition, it was selected from each individual decomposition as the ABR mode (ABR_VMD_).

The different ABR representations (ABR_narrow_, ABR_broad_, ABR_PCA_, ABR_VMD_) were finally reduced to scalar features by extracting the amplitude of the prominent wave V. Prior to that, the wave V peak latency was obtained by pooling the mean ERPs from vertex recordings across participants and conditions. Subsequently, the wave V amplitude was computed as the mean potential within a 3ms window centered at 5ms.

### Statistics

All statistical analyses to investigate effects of attention described below were based on contrasting conditions using within-participants t-tests (two–tailed) and *p*–value adjustments for multiple comparisons following the Benjamini–Hochberg FDR correction procedure [70].

ERP time–domain waveforms and time–frequency power and ITPC maps were analyzed in an exploratory manner. Waveforms were tested at each time point in [0, 250]ms and *p*-values were FDR-corrected across time (*q* = 0.05). Similarly, power and ITPC maps were analyzed at each time–frequency point within the same time interval and a frequency range of 4–512Hz and *p*-values were FDR-corrected across the spectrotemporal plane (*q* = 0.05). Due to the redundancy of the continuous wavelet transform and to obtain a more spectrotemporally localized representation of the different neural activity components contributing to the differences between conditions, FDR correction was also repeated with a more stringent level (*q* = 0.01). Furthermore, statistical analysis of ITPC comprised an additional stage at which effects were only classified as significant if they were located within a certain “area of trust”. Time–frequency points were assigned to “areas of trust” if the mean ITPC across participants indicated a significant deviation (*p* < 0.01) from a circular random distribution as analyzed via Rayleigh’s test[68].

The time–domain correlation and energy measures for ABR_narrow_, AMLR_narrow_, and ALR_narrow_ were tested in a channel–wise manner for vertex and multi–channel recordings. The resulting *p*–values for multi–channel recordings were FDR–corrected across the scalp (*q* = 0.05), separately for each full–range sub–component and each measure. Detailed statistical results for the vertex recordings are given in the SI.

Effects of attention on wave V amplitudes were individually tested for ABR_narrow_, ABR_broad_, ABR_PCA_, and ABR_VMD_. To ensure that all ABR_VMD_ representations were comparable, the peak frequencies of all selected modes were also contrasted between conditions. This was repeated for the selected ABR_PCA_ components which was complemented by comparisons of the corresponding percentages of explained variance as well as the channel–wise component weights. The resulting *p*–values for the component weights were FDR-corrected across the scalp (*q* = 0.05). The detailed significance information is provided in the SI.

## Acknowledgments

This work was partially supported by European Union (European Regional Development Fund, ERDF) and Saarland via the Center for Digital Neurotechnologies Saar (CDNS). The authors would also like to thank Simon V. Busch, Systems Neuroscience & Neurotechnology Unit, for performing the source localization analysis.

## Data & Code

The data and code can be downloaded from github after journal acceptance. Please contact the corresponding author for any interest before.

## Author Information

D.J.S., F.I.C.S., and S.A.H. designed research; F.I.C.S. performed the measurements; D.J.S., F.I.C.S, and A.M. performed research; A.M. analyzed data; D.J.S. and S.A.H. wrote the paper

## SUPPLEMENTARY MATERIAL

### The Auditory Stimuli

The auditory stimuli used in the study are shown as waveform in Fig. 5.

**Figure 5:**
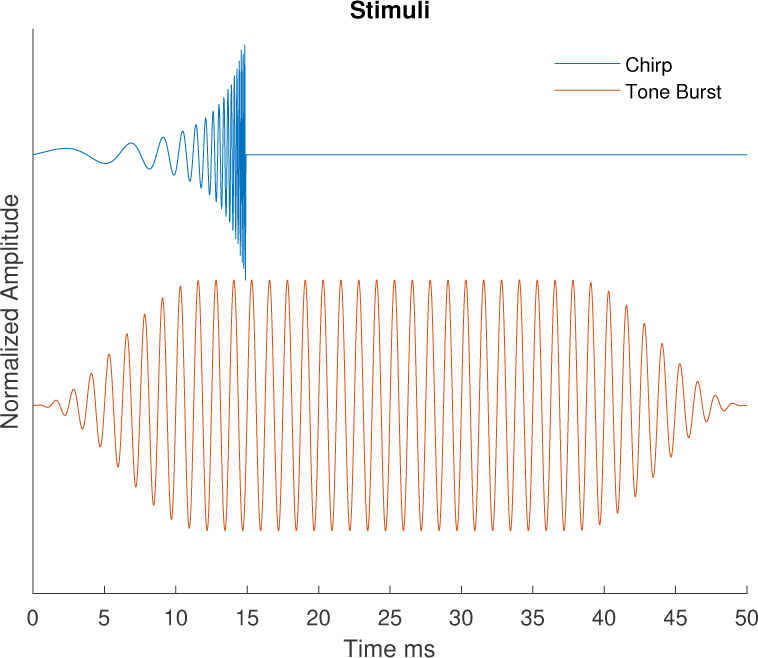
Stimuli: Chirp and tone burst with a normalized amplitude. The chirps last 14.9ms, while the tone burst is 50 ms long. The target stimuli are exactly the same, but attenuated 15dB. Note that both stimuli start with positive rising amplitude.

### Time–Frequency Analysis of the Tone ERPs

Complementary to Fig. 2 in the main text, the time–frequency analysis of full–range ERPs evoked by the tones is shown in Fig. 6.

**Figure 6:**
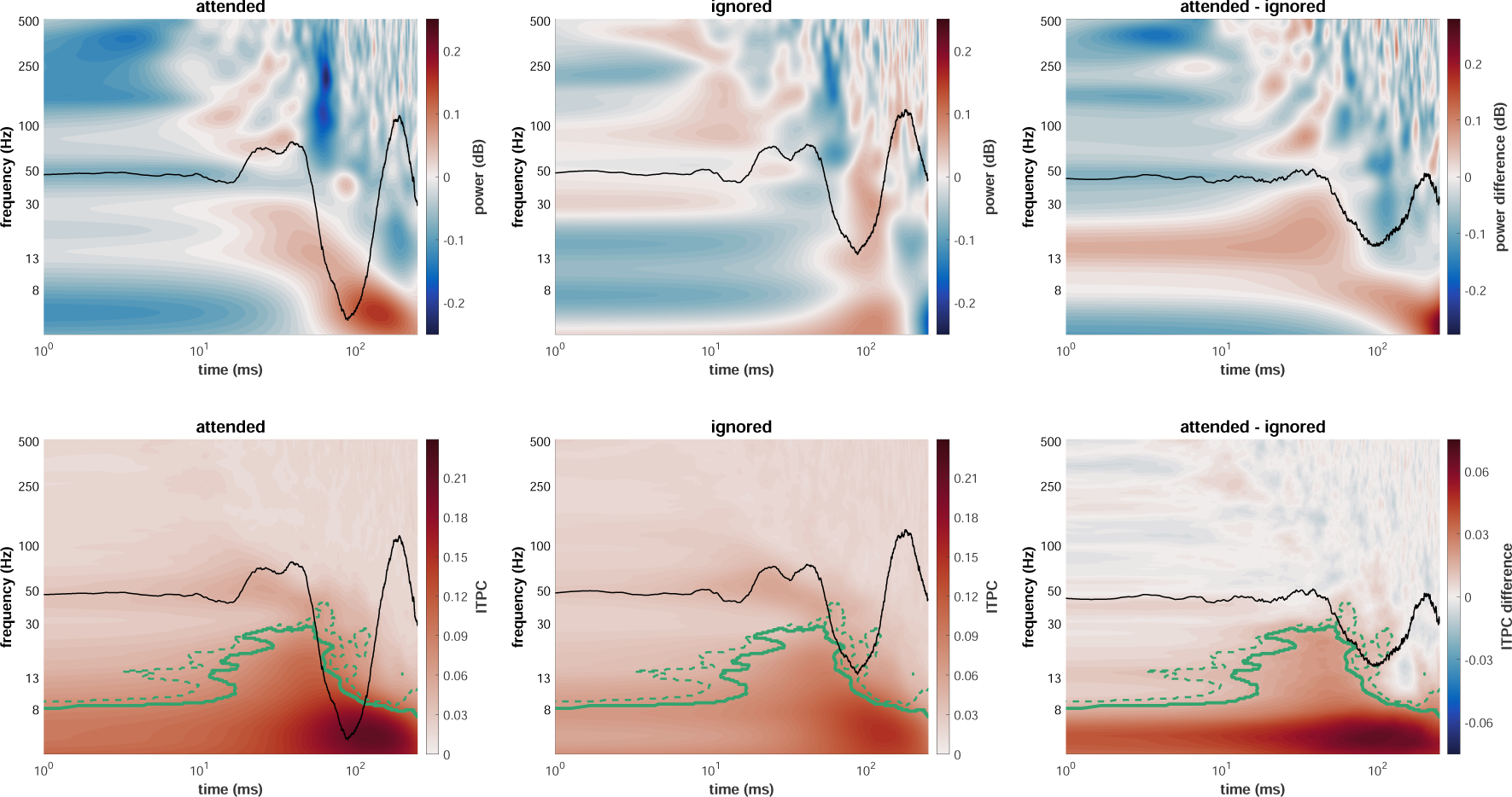
Wavelet power (top) and ITPC (bottom) analysis of vertex broadband single sweeps to attended (left) and ignored (middle) tone stimuli as well as the resulting difference map (right; attended - ignored). Black waveforms represent identically scaled condition–specific grand average ERPs (left and middle) and their difference wave (right). Time–frequency areas which indicated significant differences between conditions are represented by green lines (*p* < 0.05 for dashed lines and *p* < 0.01 for solid lines; two–tailed within–subjects t–test with FDR correction).

### Statistical Details

The wave V amplitude analyses of ABR_narrow_, ABR_broad_, ABR_PCA_, and ABR_VMD_ revealed significant effects of attention for all ABR representations. Table 1 provides a summary of the comparisons highlighted in Fig. 3 of the main text. The center frequencies of the selected modes for ABR_VMD_ were not significantly influenced by attention (*t*(30) = *−*1.71*, p* = 0.097). The same applies to the center frequencies of the selected components for ABR_PCA_ (*t*(30) = *−*1.16*, p* = 0.26) as well as their percentages of explained variance (*t*(30) = 1.33*, p* = 0.20) and weights across the scalp (all *p >* 0.05, FDR-corrected).

**Table 1:** Results of the significance analyses for the wave V amplitudes of the different ABR representations presented in Fig. 3 of the main text. All results were obtained by testing attended vs. ignored via two–tailed within–subjects t–tests.

| ABR representation | DOF | $t$-value | $p$-value |
| --- | --- | --- | --- |
| ABR<sub>narrow</sub> | 30 | 2.95 | 0.0062 |
| ABR<sub>broad</sub> | 30 | 2.47 | 0.0193 |
| ABR<sub>PCA</sub> | 30 | 2.40 | 0.0231 |
| ABR<sub>VMD</sub> | 30 | 2.68 | 0.0119 |

The time–domain correlation and energy analyses of ABR_narrow_, AMLR_narrow_, and ALR_narrow_ revealed significant effects of attention for the vast majority of channels for the correlation between the mean full–range ERP sub–components and their individual sweeps, but not for the mean energy across individual sweeps. Table 2 provides a summary of the comparisons for the vertex recordings presented in Fig. 4 (top) of the main text.

**Table 2:** Results of the significance analyses for the time–domain correlation and energy measures presented in Fig. 4 (top) of the main text. All results were obtained by testing attended vs. ignored via two–tailed within–subjects t–tests.

| ERP sub-component | measure | DOF | $t$-value | $p$-value |
| --- | --- | --- | --- | --- |
| ABR<sub>narrow</sub> | correlation | 30 | 4.19 | $2.24 \times 10^{-4}$ |
| AMLR<sub>narrow</sub> | correlation | 30 | 4.31 | $1.62 \times 10^{-4}$ |
| ALR<sub>narrow</sub> | correlation | 30 | 6.23 | $5.72 \times 10^{-7}$ |
| ABR<sub>narrow</sub> | energy | 30 | -0.27 | 0.79 |
| AMLR<sub>narrow</sub> | energy | 30 | -0.72 | 0.48 |
| ALR<sub>narrow</sub> | energy | 30 | 1.45 | 0.16 |

### Up–Chirp Behavior of Full–Range ERPs

Let us model the instantaneous frequency of a full–range ERP by 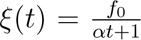 and let *γ*(*t*) be an amplitude modulation function in *t ∈* [0, 250ms] representing ABR, AMLR, and ALR amplitude differences. Then the chirp ERP can be defined by the following function: 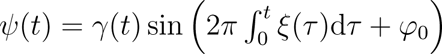. This yields

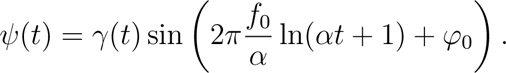

With *α* = 650, *f*_0_ = 500Hz, and *φ*_0_ = *−π* we obtain a positive deflection at 100ms in the *θ* frequency band resembling the N1 wave and the full–range ERP frequency behavior, see [32, 33]. The rapid decay of *ξ*(*t*) with 1/*t* is also reflected in Fig. 2 in the main text, showing tendentially a linear decrease of the ERP frequency and a uniform oscillation in Fig. 1, both on a logarithmic time scale; even though concurrently occurring frequency components show that the ERP is a composite signal of a larger spectrotemporal complexity than a simple up–chirp. However, ERPs reflect the general 1/*t* trend that might stem from different neuronal generators, spikes for ABRs [36, 41] and synaptic activity of pyramidal neurons for ALRs.

### Time–Frequency Super–Resolution

The resolution of frequencies around 100 *−* 200Hz on a millisecond–scale is challenging because of associated Heisenberg boxes (see, e.g., [71, 72]). This holds for Fig. 2 (main text). As alternative to, e.g., Cohen’s class time–frequency distributions (see, e.g., [72]), time–frequency super–resolution has recently been suggested to deal with these type of resolution problems [73]. Super–resolution is achieved by combining the multi–analysis of a set of (the same type) wavelets with a different support in terms of their cycles geometrically, called superlets in [73]. This combined analysis improves the frequency resolution for higher frequencies, while maintaining a good compact support in time. In Fig. 7, we have shown the grand average of the time–frequency power of the mean ABRs with super–resolution using (fractional) multiplicative superlets [73] for the attended and the ignored chirp stimulation (analytic Morlet wavelet as kernel, start cycle 2, order 2, frequencies between 4–256Hz in steps of 0.2Hz). The improved time–frequency resolution in terms of the covered Heisenberg area allows us to detangle time–frequency energy blobs around wave V. It is noticeable wave V produces an individual time– frequency blob when using super–resolution and is not the result of time–frequency ‘smearing’ due to Heisenberg’s uncertainty principle from neighboring components. The baseline normalization (here [*−*200*−−*50]ms) procedure and the statistical analysis were analog to the analysis of wavelet power for full–range ERPs.

**Figure 7:**
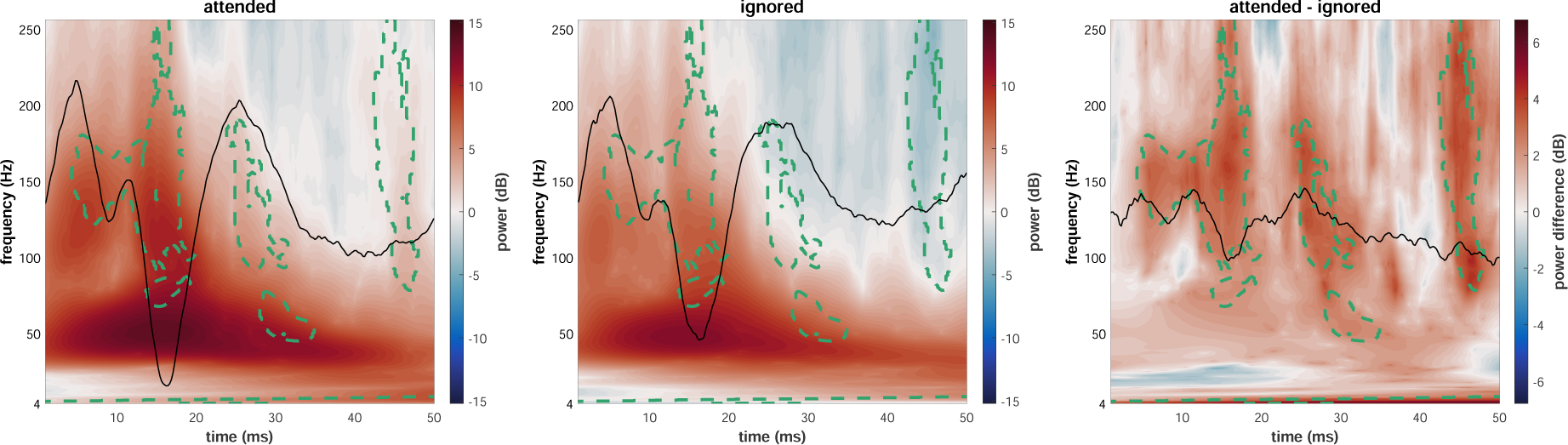
Time–frequency super–resolution power analysis within the ABR interval of mean vertex ERPs to attended (left) and ignored (middle) stimuli as well as the resulting difference map (right; attended - ignored). Black waveforms represent identically scaled condition–specific grand average ERPs (left and middle) and their difference wave (right). Time–frequency areas which indicated significant differences between conditions are represented by green dashed lines (*p* < 0.05; two–tailed within–subjects t–test with FDR correction). Note the clearly localized time–frequency component synchronized with the wave V peak latency and the significant effects reaching into the ABR latency range.

### Source Localization

Source estimation of wave V generators was carried out using BESA Research 7.1 (BESA GmbH, Germany). A pair of symmetrical dipoles was used to model the underlying sources. Dipoles were fit over the time interval 3.5–6.5ms and the regularization constant was set to 0%. BESA’s realistic FEM head model template was used for modeling geometry and conductivity of the head. Grand average ERP data were used for source estimation. For the final analysis, four channels were excluded due to missing correspondences in the 10-5 system in BESA. Finally, 120 channels recorded from electrodes placed according to the 10-5 system were used for source localization.

For both conditions, source estimation located the underlying generators of wave V in the brainstem close to the inferior colliculi. The MNI coordinates for the fit dipoles were *±*3.2mm, −37.1mm, 22.5mm in the attended case (see Fig. 8) and *±*3.8mm, −27.5mm, 22.3mm in the ignored case (see Fig. 9). Dipoles explained up to 93.5% and 89.5% of the variance for attended and ignored, respectively. Using source estimation, generators of wave V could be localized close to the physiological sources (MNI coordinates: *±*5 mm, *−*35 mm, *−*11 mm) [31, 74].

**Figure 8:**
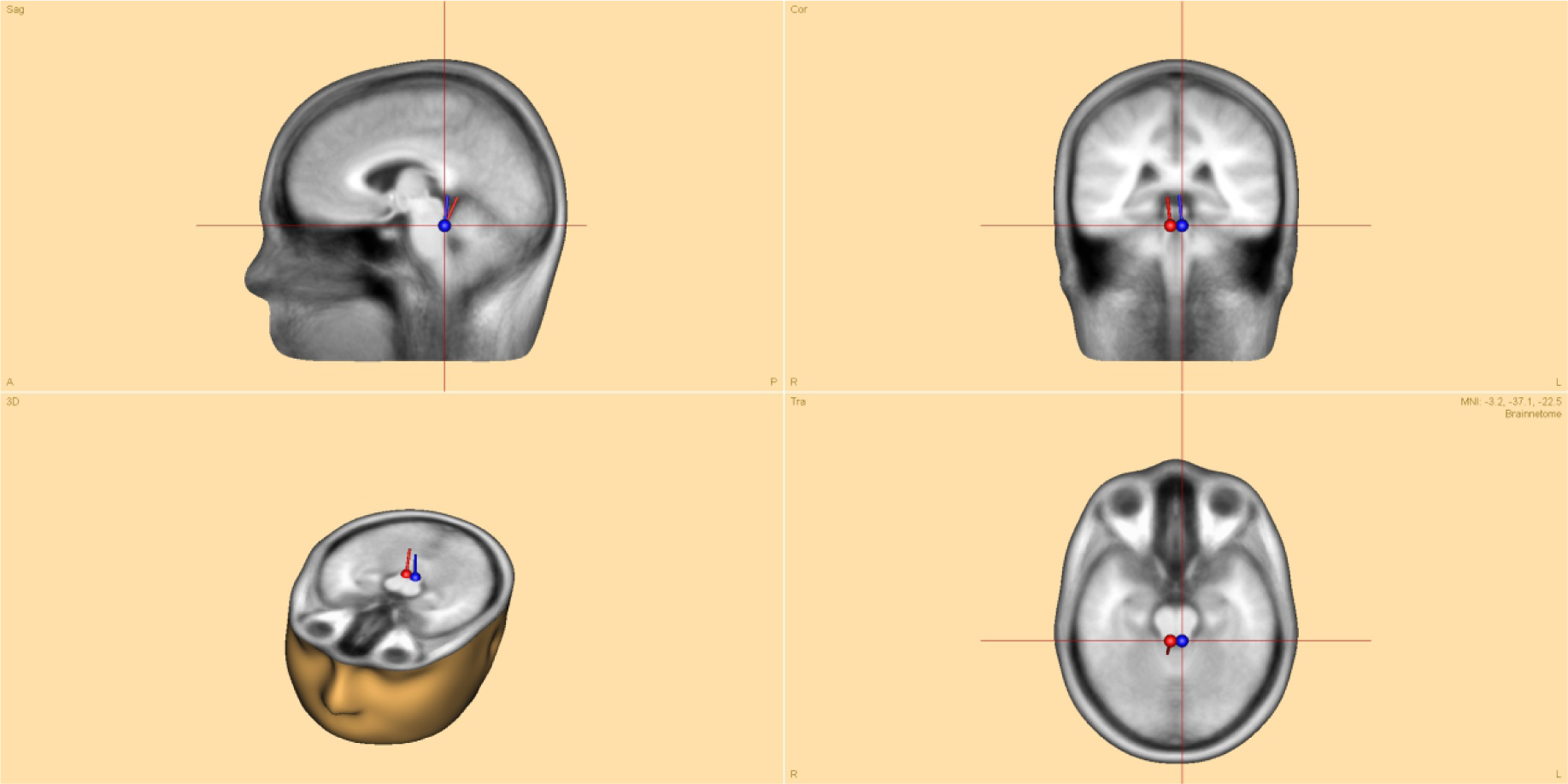
Source estimation of wave V for the attended condition, visualized in different planes. A pair of symmetrical dipoles was used to model the underlying sources over the interval from 3.5 ms to 6.5 ms. The resulting dipoles were located in the brainstem, close to the inferior colliculi.

**Figure 9:**
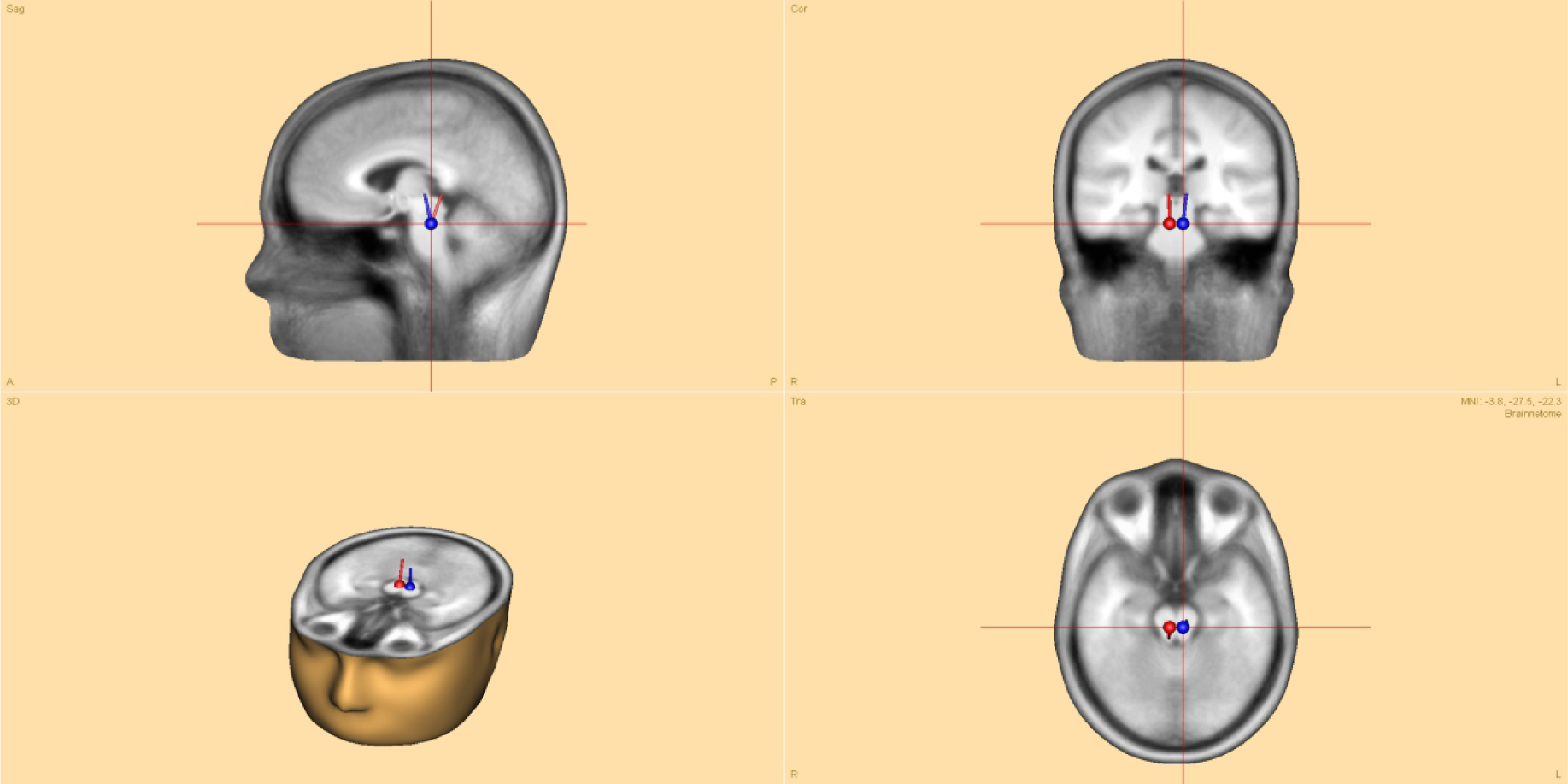
Source estimation of wave V for the ignored condition, visualized in different planes. A pair of symmetrical dipoles was used to model the underlying sources over the interval from 3.5 ms to 6.5 ms. The resulting dipoles were located in the brainstem, close to the inferior colliculi.

## References

[1] R. Näätänen. Selective attention and evoked potentials. Annals Academiac Scientiarum Fennicae, B 151:1–226, 1967.

[2] S. A. Hillyard, R. F. Hink, V. L. Schwent, and T. W. Picton. Electrical signs of selective attention in the human brain. Science, 182:177–180, 1973.

[3] R. Näätänen, A. W. Gaillard, and S. Mäntysalo. Early selective-attention effect on evoked potential reinterpreted. Acta Psychologica, 42:313–329, 1978.

[4] J. C. Hansen and S. A. Hillyard. Endogenous brain potentials associated with selective auditory attention. Electroencephalography and Clinical Neurophysiology, 49:277–290, 1980.

[5] M. H. Giard, A. Fort, Y. Mouchetant-Rostaing, and J. Pernier. Neurophysiological mechanisms of auditory selective attention in humans. Frontiers in Bioscience, 5: 84–94, 2000.

[6] M. Scherg and D. Von Cramon. Two bilateral sources of the late AEP as identified by a spatio-temporal dipole model. Electroencephalography and Clinical Neurophysiology, 62:32–44, 1985.

[7] R. Näätänen and T. W. Picton. The N1 wave of the human electric and magnetic response to sound: a review & analysis of the component structure. Psychophysiology, 24:375–425, 1987.

[8] M. G. Woldorff, C. C. Gallen, S. A. Hampson, S. A. Hillyard, C. Pantev, D. Sobel, and F. E. Bloom. Modulation of early sensory processing in human auditory cortex during auditory selective attention. Proceedings of the National Academy of Sciences, 90:8722–8726, 1993.

[9] M. G. Woldorff and S. A. Hillyard. Modulation of early auditory processing during selective listening to rapidly presented tones. Electroencephalography and Clinical Neurophysiology, 79:170–191, 1991.

[10] J. A. Winer, D. T. Larue, J. J. Diehl, and B. J. Hefti. Auditory cortical projections to the cat inferior colliculus. Journal of Comparative Neurology, 400:147–174, 1998.

[11] N. Suga, Z. Xiao, X. Ma, and W. Ji. Plasticity and corticofugal modulation for hearing in adult animals. Neuron, 36:9–18, 2002.

[12] J. M. Blackwell, A. M. Lesicko, W. Rao, M. De Biasi, and M. N. Geffen. Auditory cortex shapes sound responses in the inferior colliculus. eLife, 9:e51890, 2020.

[13] T. W. Picton, S. A. Hillyard, R. Galambos, and M. Schiff. Human auditory attention: a central or peripheral process? Science, 173:351–354, 1971.

[14] T. W. Picton, S. A. Hillyard, H. I. Krausz, and R. Galambos. Human auditory evoked potentials. Audiology and Neurootology, 36:179–190, 1974.

[15] D. L. Woods and S. A. Hillyard. Attention at the cocktail party: Brainstem evoked potentials reveal no peripheral gating. In D. A. Otto, editor, Multidisciplinary perspectives in event-related brain potential research, pages 230–233, 1978.

[16] S. A. Hackley, M. Woldorff, and S. A. Hillyard. Combined use of microreflexes and event-related brain potentials as measures of auditory selective attention. Psychophysiology, 24:632–647, 1987.

[17] S. A. Hackley, M. Woldorff, and S. A. Hillyard. Cross-modal selective attention effects on retinal, myogenis, brainstem, and cerebral evoked potentials. Psychophysiology, 27:195–208, 1990.

[18] L. Collet and R. Duclaux. Auditory brainstem evoked responses and attention. Contribution to a controversial subject. Acta Otolaryngologica, 101:439–441, 1986.

[19] J. F. Connolly, K. Aubry, N. Mcgillivary, and D. W. Scott. Human brainstem auditory evoked potentials fail to provide evidence of efferent modulation of auditory input during attentional tasks. Psychophysiology, 26:292–303, 1989.

[20] S. D. Gregory, J. A. Heath, and M. E. Rosenberg. Does selective attention influence the brain-stem auditory evoked potential? Electroencephalography and Clinical Neurophysiology, 73:557–560, 1989.

[21] T. N. Hirschhorn and P. T. Michie. Brainstem auditory evoked potentials (BAEPs) and selective attention revisited. Psychophysiology, 27:495–512, 1990.

[22] G. C. Galbraith and C. Arroyo. Selective attention and brainstem frequency-following responses. Biological Psychology, 37:3–22, 1993.

[23] A. Lehmann and M. Schönwiesner. Selective attention modulates human auditory brainstem responses: relative contributions of frequency and spatial cues. PLoS One, 9:e85442, 2014.

[24] L. Varghese, H. M. Bharadwaj, and B. G. Shinn-Cunningham. Evidence against attentional state modulating scalp-recorded auditory brainstem steady-state responses. Brain Research, 1626:146–164, 2015.

[25] A. E. Forte, O. Etard, and T. Reichenbach. The human auditory brainstem response to running speech reveals a subcortical mechanism for selective attention. eLife, 6:e27203, 2017.

[26] C. N. Price and G. M. Bidelman. Attention reinforces human corticofugal system to aid speech perception in noise. NeuroImage, 235:118014, 2021.

[27] O. Fobel and T. Dau. Searching for the optimal stimulus eliciting auditory brain-stem responses in humans. Journal of the Acoustical Society of America, 116: 2213–2222, 2004.

[28] F. I. Corona-Strauss, W. Delb, B. Schick, and D. J. Strauss. Phase stability analysis of chirp evoked auditory brainstem responses by gabor frame operators. IEEE Transactions on Neural Systems and Rehabilitation Engineering, 17:530–536, 2009.

[29] M. G. Woldorff, S. A. Hackley, and S. A. Hillyard. The effects of channel-selective attention on the mismatch negativity wave elicited by deviant tones. Psychophysiology, 28:30–42, 1991.

[30] A. R. Møller and P. J. Jannetta. Interpretation of brainstem auditory evoked potentials: results from intracranial recordings in humans. Scandinavian Audiology, 12:125–133, 1983.

[31] N. A. Parks, M. R. Hilimire, and P. M. Corballis. Visual perceptual load modulates an auditory microreflex. Psychophysiology, 46:498–501, 2009.

[32] T. W. Picton. Human Auditory Evoked Potentials. Plural Publishing Inc, San Diego, CA, USA, 2010.

[33] M. C. Kohl, E. Schebsdat, E. N. Schneider, A. Niehl, D. J. Strauss, Ö. Özdamar, and J. Bohórquez. Fast acquisition of full-range auditory event-related potentials using an interleaved deconvolution approach. Journal of the Acoustical Society of America, 145:540, 2019.

[34] C. Trenado, L. Haab, and D. J. Strauss. Corticothalamic feedback dynamics for neural correlates of auditory selective attention. IEEE Transactions on Neural Systems and Rehabilitation Engineering, 17:46–52, 2009.

[35] Y. F. Low and D. J. Strauss. A performance study of the wavelet-phase stability (WPS) in auditory selective attention. Brain Research Bulletin, 86:110–117, 2011.

[36] A. P. Rudell. A fiber tract model of auditory brain-stem responses. Electroencephalography and Clinical Neurophysiology, 67:53–62, 1987.

[37] F. I. Corona-Strauss, W. Delb, B. Schick, and D. J. Strauss. Phase stability analysis of chirp evoked auditory brainstem responses by gabor frame operators. IEEE Transactions on Neural Systems and Rehabilitation Engineering, 17:530–536, 2009.

[38] A. R. D. Thornton, M. Harmer, and B. A. Lavoie. Selective attention increases the temporal precision of the auditory N100 event-related potential. Hearing Research, 230:73–79, 2007.

[39] C. Trenado, L. Haab, W. Reith, and D. J. Strauss. Biocybernetics of attention in the tinnitus decompensation: an integrative multiscale modeling approach. Journal of Neuroscience Methods, 178:237–247, 2009.

[40] D. J. Strauss. Verfahren und Anordnung zur objektiven Kontrolle der Messqualität und zur Erkennung von ereigniskorrelierten Potenzialen in den Ableitungen einer neuronalen Aktivität, 2016. Patent DE102011114045.

[41] A. R. Møller and P. J. Jannetta. Neural generators of the auditory brainstem response. In J. T. Jacobson, editor, The auditory brainstem response, pages 13–31, 1985.

[42] B. Yvert, C. Fischer, M. Guénot, P. Krolak-Salmon, J. Isnard, and J. Pernier. Simultaneous intracerebral EEG recordings of early auditory thalamic and cortical activity in human. European Journal of Neuroscience, 16:1146–1150, 2002.

[43] T. Rinne, M. H. Balk, S. Koistinen, T. Autti, K. Alho, and M. Sams. Auditory selective attention modulates activation of human inferior colliculus. Journal of Neurophysiology, 100:3323–3327, 2008.

[44] A. Schüller, A. Schilling, P. Krauss, S. Rampp, and T. Reichenbach. Attentional modulation of the cortical contribution to the frequency-following response evoked by continuous speech. Journal of Neuroscience, 43:7429–7440, 2023.

[45] J. Benhamou, M. Le Van Quyen, and G. Marrelec. Time-frequency analysis of event-related brain recordings: Connecting power of evoked potential and intertrial coherence. IEEE Transactions on Biomedical Engineering, 70:1599–1610, 2023.

[46] Z. Mortezapouraghdam, F. I. Corona-Strauss, T. Takahashi, and D. J. Strauss. Reducing the effect of spurious phase variations in neural oscillatory signals. Frontiers in Computational Neuroscience, 12:82, 2018.

[47] G. Buzsáki, C. Anastassiou, and C. Koch. The origin of extracellular fields and currents — EEG, ECoG, LFP and spikes. Nature Reviews Neuroscience, 13:407–420, 2012.

[48] M. Rosenblum, A. Pikovsky, J. Kurths, C. Schäfer, and P. A. Tass. Phase synchronization: from theory to data analysis. In A. J. Hoff, editor, Handbook of Biological Physics, pages 279–321, 2001.

[49] P. van der Wel and H. van Steenbergen. Pupil dilation as an index of effort in cognitive control tasks: A review. Psychonomic Bulletin & Review, 25:2005–2015, 2018.

[50] R. Desimone and J. Duncan. Neural mechanisms of selective visual attention. Annual Review of Neuroscience, 18:193–222, 1995.

[51] T. Womelsdorf and P. Fries. The role of neural synchronization in selective attention. Current Opinion in Neurobiology, 17:154–160, 2007.

[52] G. G. Gregoriou, S. J. Gotts, H. Zhou, and R. Desimone. High-frequency, long-range coupling between prefrontal and visual cortex during attention. Science, 324: 1207–1210, 2009.

[53] G. Buzsáki. The Brain from Inside Out. Oxford University Press Inc, New York, NY, USA, 2021.

[54] Buschman T. J. and Kastner S. From behavior to neural dynamics: An integrated theory of attention. Neuron, 88:127–144, 2015.

[55] H. M. Oberle, A. N. Ford, J. E. Czarny, M. M. Rogalla, and P. F. Apostolides. Recurrent circuits amplify corticofugal signals and drive feedforward inhibition in the inferior colliculus. Journal of Neuroscience, 43:5642–5655, 2023.

[56] B. M. Sayers, H. A. Beagley, and W. R. Henshall. The mechanism of auditory evoked EEG responses. Nature, 247:481–483, 1974.

[57] B.-K. Min, N. A. Busch, S. Debener, C. Kranczioch, S. Hanslmayr, A. K. Engel, and C. S. Herrmann. The best of both worlds: Phase–reset of human EEG alpha activity and additive power contribute to ERP generation. International Journal of Psychophysiology, 65:58–68, 2007.

[58] A. P. Burgess. Towards a unified understanding of event-related changes in the EEG: The firefly model of synchronization through cross-frequency phase modulation. PLoS ONE, 7:e45630, 2012.

[59] A. A. Studenova, A. Villringer, and V. V. Nikulin. Non-zero mean alpha oscillations revealed with computational model and empirical data. PLOS Computational Biology, 18:e1010272, 2022.

[60] B. Telenczuk, V. V. Nikulin, and G. Curio. Role of neuronal synchrony in the generation of evoked EEG/MEG responses. Journal of Neurophysiology, 104:3557–3567, 2010.

[61] E. C. Wilson, G. Schlaug, and C. Pantev. Listening to filtered music as a treatment option for tinnitus: A review. Music Perception, 27:327–330, 2010.

[62] O. Wegner and T. Dau. Frequency specificity of chirp-evoked auditory brainstem responses. Journal of the Acoustical Society of America, 111:1318–1329, 2002.

[63] D. J. Strauss, F. I. Corona-Strauss, A. Schroeer, P. Flotho, R. Hannemann, and S. A. Hackley. Vestigial auriculomotor activity indicates the direction of auditory attention in humans. eLife, 9:e54536, 2020.

[64] D. Chicco and G. Jurman. The advantages of the matthews correlation coefficient (MCC) over F1 score and accuracy in binary classification evaluation. BMC Genomics, 21:6, 2020.

[65] A. Delorme and S. Makeig. EEGLAB: an open source toolbox for analysis of single-trial EEG dynamics including independent component analysis. Journal of Neuroscience Methods, 134:9–21, 2004.

[66] J. M. Lilly and S. C. Olhede. Generalized morse wavelets as a superfamily of analytic wavelets. IEEE Transactions on Signal Processing, 60:6036–6041, 2012.

[67] S. Morales and M. E. Bowers. Time-frequency analysis methods and their application in developmental EEG data. Developmental Cognitive Neuroscience, 54: 101067, 2022.

[68] S. Rao Jammalamadaka and A. Sengupta. Topics in Circular Statistics. World Scientific Pub Co Inc, 2001.

[69] K. Dragomiretskiy and D. Zosso. Variational mode decomposition. IEEE Transactions on Signal Processing, 62:531–544, 2014.

[70] Y. Benjamini and Y. Hochberg. Controlling the false discovery rate: A practical and powerful approach to multiple testing. Journal of the Royal Statistical Society: Series B (Methodological), 57:289–300, 1995.

[71] I. Daubechies. Ten Lectures on Wavelets. SIAM, Philadelphia, PA, USA, 1992.

[72] S. Mallat. A Wavelet Tour of Signal Processing. Academic Press, San Diego, CA, USA, second edition, 1999.

[73] V. V. Moca, H. Bârzan, A. Nagy-Dăbâcan, and R. C. Muresan. Time-frequency super-resolution with superlets. Nature Communications, 12:337, 2021.

[74] S. Da Costa, J. Clément, R. Gruetter, and Ö. Ipek. Evaluation of the whole auditory pathway using high-resolution and functional MRI at 7T parallel-transmit. PLoS ONE, 16:e0254378, 2021.

